# EllipTrack: A Global-Local Cell-Tracking Pipeline for 2D Fluorescence Time-Lapse Microscopy

**DOI:** 10.1101/2020.04.13.036756

**Authors:** Chengzhe Tian, Chen Yang, Sabrina L. Spencer

**Affiliations:** Department of Biochemistry; Department of Molecular, Cellular, and Developmental Biology; BioFrontiers Institute, University of Colorado Boulder, Boulder CO 80303, USA

## Abstract

Time-lapse microscopy provides an unprecedented opportunity to monitor single-cell dynamics. However, tracking cells for long periods of time remains a technical challenge, especially for multi-day, large-scale movies with rapid cell migration, high cell density, and drug treatments that alter cell morphology/behavior. Here, we present EllipTrack, a global-local cell-tracking pipeline optimized for tracking such movies. EllipTrack first implements a global track-linking algorithm to construct tracks that maximize the probability of cell lineages, and then corrects tracking mistakes with a local track-correction module where tracks generated by the global algorithm are systematically examined and amended if a more probable alternative can be found. Through benchmarking, we show that EllipTrack outperforms state-of-the-art cell trackers and generates nearly error-free cell lineages for multiple large-scale movies. In addition, EllipTrack can adapt to time- and cell density-dependent changes in cell migration speeds, requires minimal training datasets, and provides a user-friendly interface. EllipTrack is available at github.com/tianchengzhe/EllipTrack.

## Introduction

Biological processes are highly dynamic and display varying degrees of cell-to-cell heterogeneity. Time-lapse imaging enables analysis of single-cell dynamics, providing longitudinal information that is not readily accessible by other single-cell methods. In a typical 2D fluorescence time-lapse microscopy experiment, cells are labeled with fluorescent markers for the signals of interest and imaged periodically under a microscope to generate a movie. A computational tool called a “cell tracker” is then applied to the movie to track cells over time and extract signals from each cell (Kudo et al., 2017). Due to its ability to visualize molecular activities in living cells in real time, time-lapse microscopy has been used to study a wide range of biological questions.

A classical cell tracker typically consists of three steps (Figure 1A). First, in Segmentation, cell nuclei are identified from the images using a fluorescent nuclear marker such as Histone 2B (H2B). Then, in Track Linking, nuclei are mapped between neighboring frames and biological events such as mitosis and apoptosis are identified. Finally, in Signal Extraction, signals are extracted from the regions of interest in each cell to gain biological insights. During the last decades, enormous effort has been made to improve the accuracy of each step (He et al., 2017; Hernandez et al., 2018; Maška et al., 2014; Moen et al., 2019; Ulman et al., 2017). Some cell trackers also attempt to perform Segmentation and Track Linking jointly, such that the properties of cell tracks from previous frames can be used as prior information to track later frames (Amat et al., 2014; Arbelle et al., 2018; Cappell et al., 2016). Despite these efforts, cell tracking remains a bottleneck in the field of time-lapse microscopy due to the necessity of achieving extremely high accuracy at each frame (Skylaki et al., 2016). Consequently, manual verification and correction is often required (Han et al., 2019; Hilsenbeck et al., 2016), which limits the scale of experiments and results in large amounts of data remaining unextracted and unquantified.

**Figure 1.**
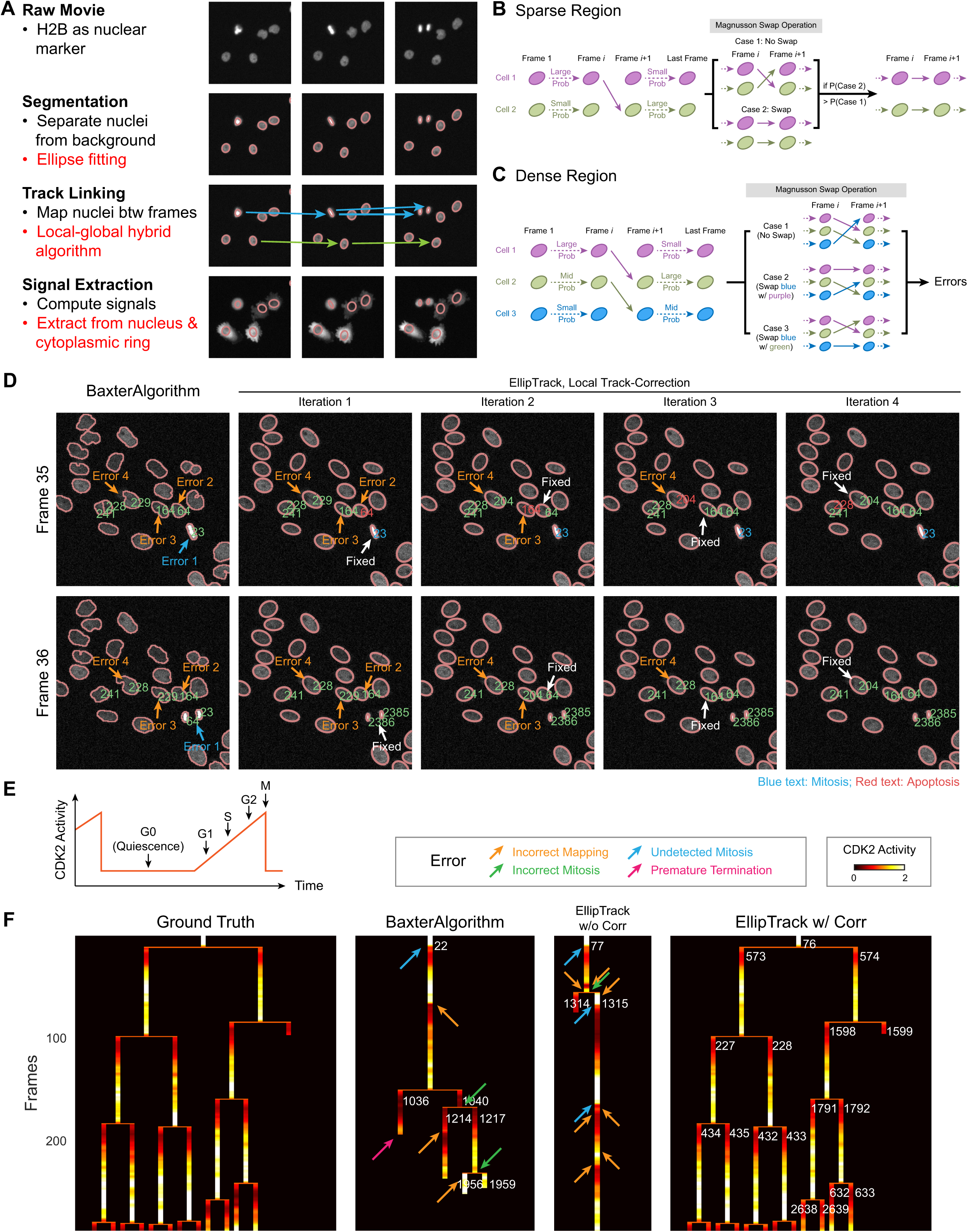
The EllipTrack pipeline. (A) Workflow of cell trackers. Red text indicates the methods implemented in EllipTrack. (B) Schematic illustration of the Magnusson swap operation. (C) Schematic illustration of the limitation of the Magnusson swap operation for cells in densely populated regions. (D) Correction of the tracking mistakes of BaxterAlgorithm by the local track-correction module of EllipTrack. Contours of segmented objects are plotted on top of the image of the nuclear marker. IDs of selected cell tracks are plotted next to their respective nuclei. Red IDs indicate apoptosis; blue IDs indicate mitosis. Tracking mistakes are indicated by the orange (incorrect mapping) or blue (undetected mitosis) arrows. Fixed errors are indicated by the white arrows at the iteration where they are fixed and removed in later iterations. Track IDs of EllipTrack are adjusted such that unmodified cell tracks always receive the same ID. For EllipTrack, Track 241 and 228 map to the same ellipse. Dataset: MCF10A-1. (E) Schematic illustration of classifying cell-cycle phases based on CDK2 activity. (F) Comparison of identified cell lineages with different cell trackers. Cell lineages are visualized in heatmaps as described previously (Wolff et al., 2018) and colored by CDK2 activity. Cell track IDs are plotted next to the births of their respective cells except in the final frames due to space constraints. Arrows indicate tracking mistakes, as explained in the legend. The “Density” option was used to infer cell migration speeds over time. The tracking result of this cell lineage is also visualized in Movie S1. Dataset: MCF10A-1.

A major breakthrough of cell tracker development in recent years was the creation of the global track-linking algorithm by Magnusson and colleagues, which aims to construct cell tracks by maximizing the overall probability of cell lineages (Magnusson et al., 2015). Here, a cell track maps a cell in every movie frame from its appearance to its disappearance, and a cell lineage is a tree of cell tracks representing mother/daughter relationships. To do so, this algorithm first uses machine learning tools to infer the probability of cell overlapping, cell migration, and other biological events (mitosis, apoptosis, etc). It then uses the Viterbi algorithm to iteratively search for and assemble the cell track that results in the greatest increase in the probability of existing cell lineages, until this probability can no longer be improved. Due to its consideration of the entire movie rather than only two neighboring frames, this algorithm has been consistently rated as a top performer in the Cell Tracking Challenge (Maška et al., 2014; Ulman et al., 2017) and has been implemented in several recent cell trackers (Cooper et al., 2017; Hernandez et al., 2018; Magnusson et al., 2015).

However, the use of Viterbi algorithm is also a major source of tracking mistakes. To illustrate this point, consider two neighboring cells between two neighboring frames (Frame *i* and *i*+1) where the first cell’s track before Frame *i* contributes a much greater increase to the overall probability of cell lineages than the second cell’s track, but its track after Frame *i*+1 contributes less (Figure 1B). Since the aim is to find the single cell track leading to the greatest increase in the overall probability, this algorithm will map the first cell in Frame *i* to the second cell in Frame *i*+1, despite the suboptimal probability between these two frames. Correspondingly, the second cell in Frame *i* will be mapped to the first cell in Frame *i*+1. To fix this tracking mistake, Magnusson *et al.* introduces a swap operation: when constructing a new cell track for cell 2, the algorithm examines whether swapping this cell track with an existing one (cell 1) will increase the overall probability of cell lineages, and will make the swap if the probability indeed increases (Figure 1B). While this operation works well for small-scale movies such as those in the Cell Tracking Challenge where cells are sparsely populated, or more precisely, where cells migrate significantly slower than the average distance between cells, it is ineffective for modern terabyte-scale movies at the forefront of the time-lapse microscopy field where cells are continuously imaged on a multi-well plate for many days. In these movies, cells undergo multiple rounds of mitoses such that the last frame contains up to thousands of cells, and this high cell density often results in multiple neighboring cells being mis-tracked by the Viterbi algorithm at the same frame. Since the swap operation only allows the newly constructed cell track to be swapped with at most one existing track, this operation can only correct a fraction of mistakes. For example, consider the scenario with three neighboring cells where the first cell in Frame *i* is mapped to the second cell in Frame *i*+1 and the second cell is mapped to the third cell (Figure 1C). When tracking the third cell, no matter which existing track it swaps with, the algorithm will only be able to fix at most one tracking mistake, and the resulting cell lineages are still erroneous (Figure 1C). Therefore, developing an optimized swap operation that corrects the mistakes in the densely populated regions is necessary.

Here, we present EllipTrack, a global-local hybrid cell-tracking pipeline. The key innovation of EllipTrack is the user-friendly packaging of the best existing algorithms combined with new local track-swapping algorithms that are executed after the Magnusson global track-linking algorithm. Our new local track-correction algorithms (hereafter local track-correction module) optimize the Magnusson swap operation and offer a significantly higher capacity to correct tracking mistakes, especially for modern multi-day, large-scale movies with densely populated cells. Through benchmarking, we show that EllipTrack outperforms current state-of-the-art methods. We also demonstrate the practical aspects of EllipTrack, such as the ability to infer time- or cell density-dependent migration speeds, the small amount of training data required, and the user-friendliness of the interfaces.

## Results

### The EllipTrack Pipeline

EllipTrack implements the traditional three-step procedure, but introduces multiple advanced features (Figure 1A, S1A). In Segmentation, EllipTrack approximates the shapes of cell nuclei as ellipses and segments cells by applying an ellipse-fitting algorithm to the nuclear contours (Zafari et al., 2015). This method shows improved performance to separate overlapping nuclei and has been implemented in other cell trackers (Türetken et al., 2016) (Figure S1B). In Track Linking, EllipTrack first constructs cell lineages with the Magnusson global track-linking algorithm (Magnusson et al., 2015), and then executes our local track-correction module to correct tracking mistakes (detailed below). Finally, in Signal Extraction, EllipTrack calculates the signals from the nuclear and cytoplasmic ring regions of each ellipse. Signals are extracted only from the pixels that do not overlap with those of other cells, in order to avoid interference from neighboring cells (Figure S1C and S1D).

The local track-correction module consists of multiple steps, each addressing a different aspect of tracking mistakes. The core step optimizes the Magnusson swap operation to fix the mistakes in densely populated regions, illustrated in Figure 1C. Due to combinatorial explosion, it is difficult to implement an algorithm where all erroneous cell tracks can be corrected simultaneously. Instead, we use an iterative strategy where tracking mistakes are progressively fixed. In each iteration, the core step examines every two cell tracks between every two neighboring frames (i.e. a local algorithm) and determines whether there exists an alternative track configuration that will significantly increase the probability of cell lineages. The alternative track configurations are chosen to correct common tracking mistakes. For example, for two cell tracks both mapping from one frame to the next (Figure S1E), EllipTrack will consider five alternatives: swapping two cell tracks as done by the Magnusson swap operation (Case 1; correct track swaps), assigning one cell track for mitosis while terminating the other one (Cases 2-3; detect missed mitosis events), and swapping two cell tracks and then breaking one of them into pieces (Cases 4-5; remove unlikely migrations). If there exist alternative configurations that increase the probability of cell lineages over a pre-defined threshold, the alternative with the greatest fold-change will replace the existing cell track configuration. Meanwhile, if no alternative meets the threshold, the existing configuration will be kept. Since we usually require the threshold fold-change to be much greater than 1, the main features of the Magnusson global track-linking algorithm will be kept and only the significant mistakes will be corrected, thus achieving a balance between the global track-linking algorithm and the local track-correction module (i.e. a global-local algorithm). This process will then repeat until no more changes to the cell lineages can be made. In addition to the core step, the local track-correction module also fixes tracking mistakes related to under-segmentation, over-segmentation, and undetected mitosis events, as detailed in STAR Methods.

### EllipTrack Identifies Nearly Error-Free Cell Lineages

To test the power of our local track-correction module, we selected a cell tracker called BaxterAlgorithm as the sample implementation of the Magnusson global track-linking algorithm (Magnusson et al., 2015), and applied it to a 48 hr sample movie of untreated MCF10A mammary epithelial cells expressing a nuclear marker (H2B) and a cell-cycle marker (CDK2 sensor) (Spencer et al., 2013). We then imported the tracking results to EllipTrack and examined whether our local track-correction module could fix the mistakes made by BaxterAlgorithm. As shown in Figure 1D, we visualized the segmentation outcomes by plotting the contours of identified nuclei on top of the image of the H2B channel. We also displayed the IDs of selected cell tracks next to the nuclei to which they mapped. Track IDs for regular cells are green, whereas a blue or red track ID indicates that the corresponding cell undergoes mitosis or apoptosis, respectively. As shown in the first column, BaxterAlgorithm incorrectly tracked multiple neighboring cells, including not detecting the mitosis event of Track 23, and mapping Track 228, 229, 164, and 64 to the incorrect neighboring cells. This is a typical example of tracking mistakes in densely populated regions where the Magnusson swap operation is ineffective. In comparison, EllipTrack can fix all these mistakes with its local track-correction module within four iterations (Figure 1D, Column 2-5). In the first iteration, the mitosis event of Track 23 was correctly detected, and Track 64 was terminated at Frame 35. Next, Track 164 and 64 were swapped at Frame 35 such that Track 64 was now correctly tracked and Track 164 was terminated. This process was repeated in the last two iterations, resulting in all relevant cells being correctly tracked. Therefore, our local track-correction module is capable of correcting mistakes that were overlooked by BaxterAlgorithm.

We next examined the ability of the entire EllipTrack pipeline to identify error-free cell lineages. To this end, we applied EllipTrack (both with and without local track-correction) and BaxterAlgorithm to the previously mentioned MCF10A movie and visualized the identified cell lineages on signal-based heatmaps. Here, we selected CDK2 activity as the signal of interest, where CDK2 activity is calculated as the ratio of the cytoplasmic ring signal relative to the nuclear signal of the DHB-based sensor (Spencer et al., 2013) (Figure S1F). This sensor is a widely used indicator of cell-cycle progression, as CDK2 activity is absent in quiescent (G0) cells and gradually increases throughout G1, S, and G2 phases of the cell cycle. Mitosis can be marked by a rapid drop in CDK2 activity (Spencer et al., 2013) (Figure 1E). It is therefore possible to evaluate the quality of identified cell lineages by examining whether they match the expected CDK2 activity. As shown in Figure 1F, with tracking results of this lineage visualized in Movie S1, both BaxterAlgorithm and EllipTrack before local track correction made numerous mistakes. For example, both trackers mapped cell tracks to the incorrect neighboring cells (orange arrow), as visualized by an abrupt change in CDK2 activity without mitosis, and assigned multiple incorrect mitosis events (green arrow), as visualized by a split of cell lineages without the characteristic rise and fall of CDK2 activity. Consequently, no cells in this lineage were correctly tracked throughout the entire movie. In comparison, the cell lineage identified by EllipTrack after local track correction matched the ground truth established via manual tracking, and all cells in this lineage were correctly tracked.

Finally, we performed a systematic benchmark of EllipTrack against BaxterAlgorithm and two other popular cell trackers: iLastik (Berg et al., 2019; Sommer et al., 2011) and a cell tracker from the Meyer Lab first used in (Cappell et al., 2016) and made publicly available in (Cappell et al., 2018) (Table S1). We evaluated their performance on six different movies from multiple sources: two 46 hr movies (92 frames) of untreated HeLa cells from the Cell Tracking Challenge (Maška et al., 2014; Ulman et al., 2017), one 21 hr movie (107 frames) of BJ5TA cells treated with control siRNA from the Meyer Lab (Cappell et al., 2016), one 100 hr movie (400 frames) of untreated RPE-hTERT cells from the Lahav Lab (Arbelle et al., 2018), the 48 hr movie (288 frames) of untreated MCF10A cells mentioned previously, and a 120 hr movie (482 frames) of A375 melanoma cells treated with 1 μM dabrafenib at the 24 hr mark from our lab (Table S2). These movies cover a broad spectrum of biological (e.g. cell lines, drug-treatment conditions) and technical parameters (e.g. movie lengths, cell densities, imaging protocols). Three experiments filmed cancer cells (HeLa and A375) and one experiment filmed cells under drug treatment (A375). Cells in these movies have irregular morphology and behaviors. Furthermore, three movies (MCF10A, A375, and RPE-hTERT) are multi-day, large-scale movies with up to 5 generations of cells, and have much higher complexity compared to the common datasets benchmarked in the Cell Tracking Challenge (Table S2). Thus, these movies well-represent current real-world cell tracking problems. We benchmarked the cell trackers on six criteria (STAR Methods). Two criteria were adopted from the Cell Tracking Challenge: SEG, a measure of <u>seg</u>mentation quality based on the pixel-match (Jaccard similarity index) between segmented nuclei and manual annotation; and TRA, a measure of <u>tra</u>cking accuracy based on the similarity (normalized Acyclic Oriented Graph Matching measure) between identified cell lineages and manual annotation. We also include four criteria from the perspective of biologists: %CORR_S, the percentage of cell nuclei that were <u>corr</u>ectly <u>s</u>egmented before Track Linking, as evaluated by manual inspection; #COMP, the number of <u>comp</u>lete tracks where cells were continuously tracked throughout the movie; #MIS_T, the average number of <u>mis</u>takes among the complete tracks; and %CORR_T, the percentage of complete tracks that were <u>corr</u>ectly tracked.

Benchmarked results are summarized in Table 1. For Segmentation, EllipTrack did not achieve a high pixel-match with manual annotation (low SEG score) due to the approximate nature of ellipse fitting, though it correctly segmented most cells in the movies as evidenced by top scores for %CORR_S. This result suggests that EllipTrack is suitable for biological research where cells are hard to identify and the identification of perfect nuclear contours is not absolutely necessary. For Track Linking, benchmarking shows that the local track-correction module of EllipTrack corrected a substantial number of tracking mistakes, such that the tracking results after correction achieved one of the top track linking scores across almost all movies. The improvement was especially significant for the three large-scale datasets where cells were more crowded and more tracking mistakes were made by BaxterAlgorithm. Other cell trackers may have higher scores in some criteria. For example, for the MCF10A movie (“MCF10A-1”), Cappell *et al.* tracked very few cells but with high accuracy, while BaxterAlgorithm and iLastik tracked more cells but with a much higher error rate. However, the former would correspondingly have a very limited coverage of the diverse cell behaviors in the movie, while the latter two would require significant manual effort to correct tracking mistakes before downstream analysis could be performed. In comparison, EllipTrack achieved a proper balance between coverage and accuracy, as it obtained more than 70% of all possible complete tracks, and more than half of identified complete tracks were error-free. Together, this benchmarking effort indicates that EllipTrack outperformed the other state-of-the-art cell trackers and demonstrates the power of the local track-correction module to obtain nearly error-free cell lineages.

**Table 1.**
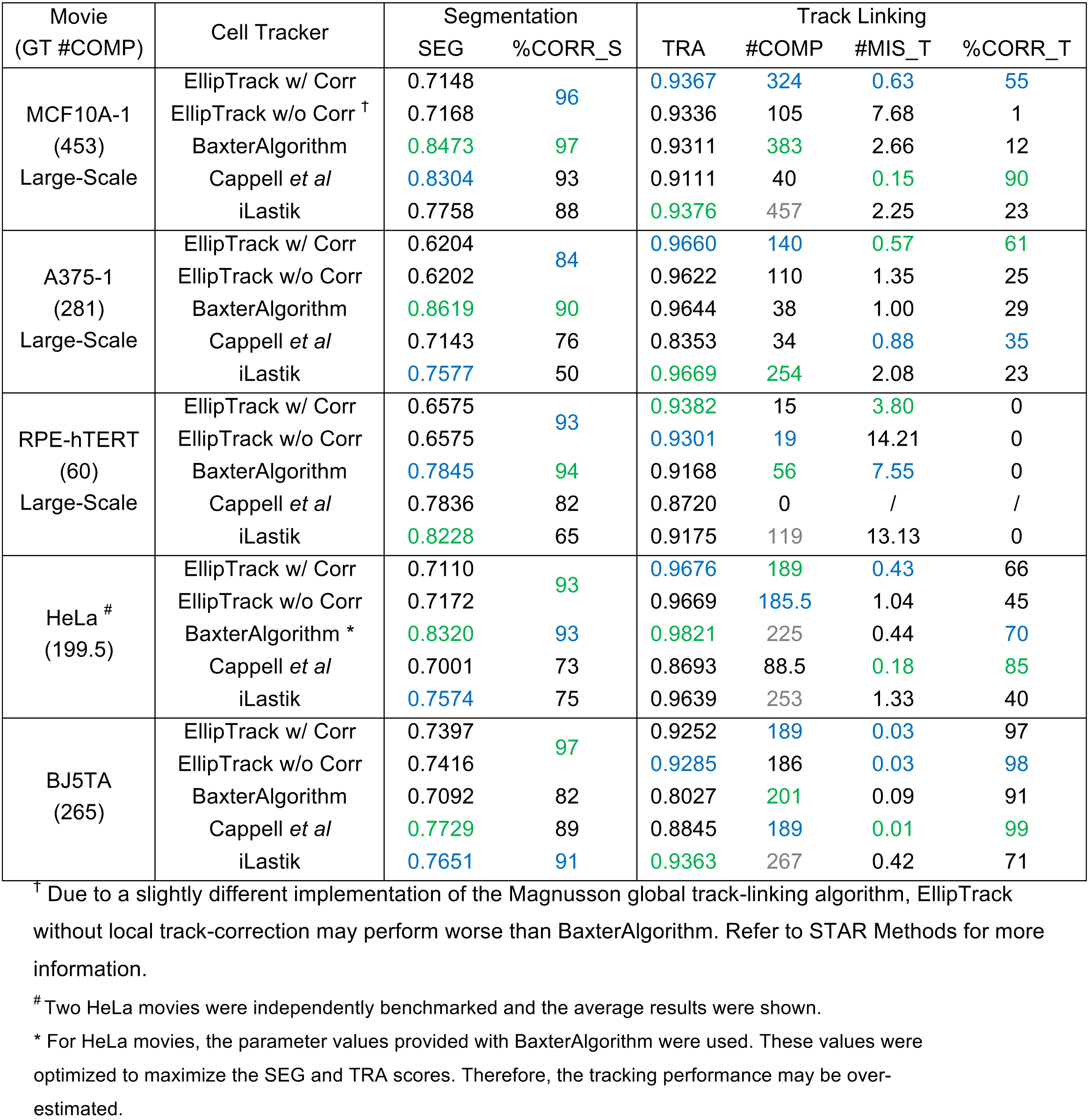
Results of cell tracker benchmark. A good tracking performance is marked by a low #MIS_T score and high scores for other criteria. For each criterion, the best and second-best scores are highlighted in green and blue, respectively. The three multi-day, large-scales movies are indicated with text “Large-Scale” under the “Movie” column. The ground truth number of complete tracks (GT #COMP) was extracted from the manually annotated cell lineages and were shown under the “Movie” column. #COMP scores greater than the ground truth value were colored in gray, as the extra complete tracks must contain tracking mistakes.

### EllipTrack Can Infer Time- and Density-Dependent Cell Migration Speed

Cells often adjust their migration speed in response to environmental changes, such as addition of drugs or changes in the local crowdedness (Ridley et al., 2003). For example, examination of a typical cell in the previously mentioned MCF10A movie revealed a clear inverse relationship between the migration speed (# pixels traveled per frame) and the number of cells in the local neighborhood (Figure 2A). However, current cell trackers implementing the Magnusson global track-linking algorithm, including BaxterAlgorithm, assume a constant migration speed over the entire movie. When tracking the cell in Figure 2A, these cell trackers are likely to terminate the cell track prematurely and re-initiate a new one in the next frame in the region of low cell density, as the probability for track termination and re-initiation can be higher than the probability for migration with an underestimated speed. Meanwhile, when the local density is high, these trackers may map the cell track to an incorrect neighboring cell. Consequently, the identified cell lineages will contain numerous mistakes.

**Figure 2.**
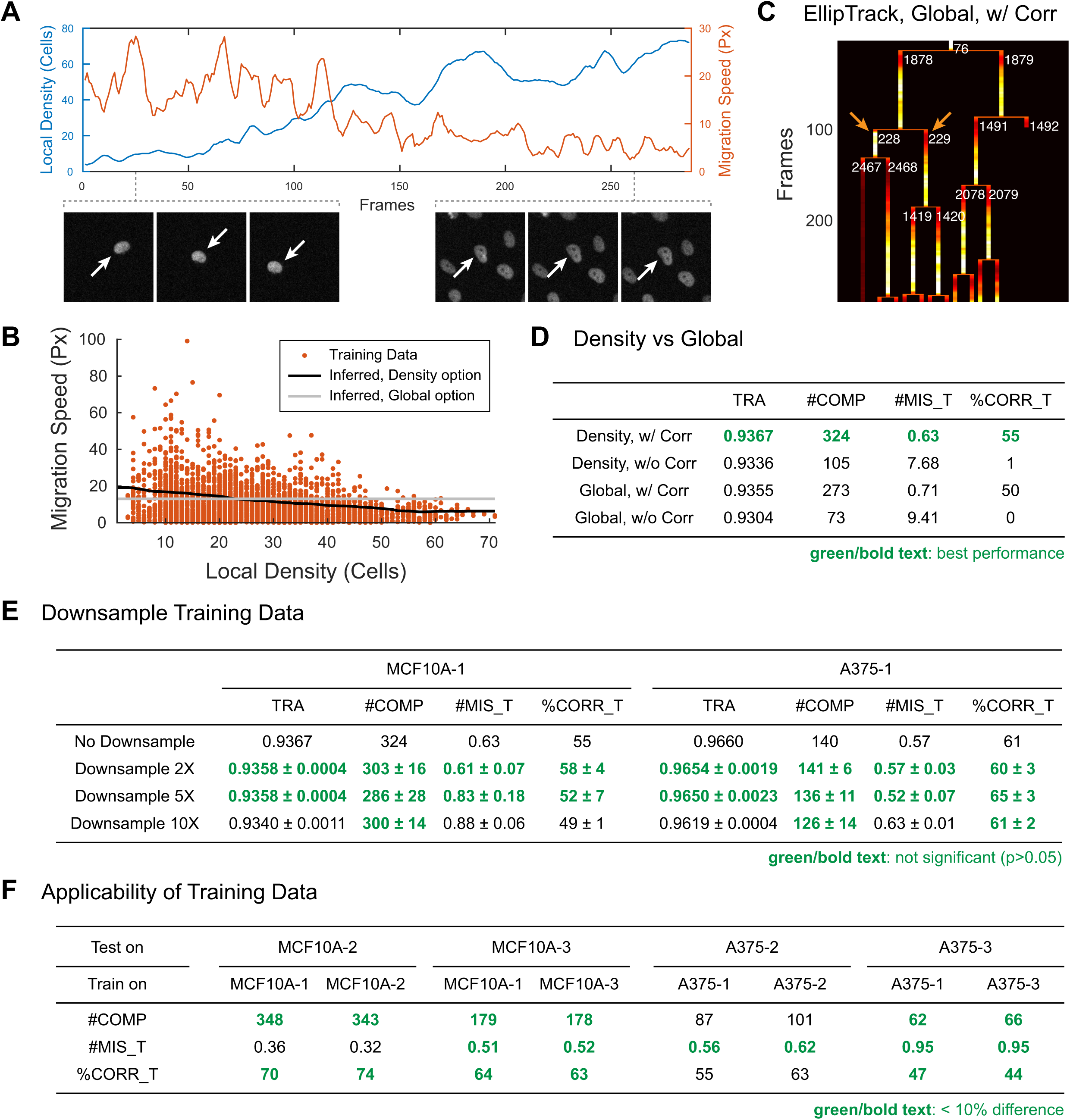
Inference of cell migration speeds and characterization of training datasets. (A) Cell migration speeds are density-dependent. Top: comparison of migration speed (Euclidean distance per frame, in pixels) and number of cells in the local neighborhood at each frame for a single cell. Bottom: images of the cell at the sparsely (left) and densely (right) populated regions. Images from three consecutive frames are shown. White arrows indicate the cell plotted in the top panel. Dataset: MCF10A-1. (B) Inference of cell migration speed from training datasets. Dots represent the migration of cells from one frame to the next; one dot is plotted for the number of pixels travelled in the x-direction, and a second dot is plotted for the number of pixels travelled in the y-direction. Black and gray lines indicate the inferred cell migration speeds with the “Density” and “Global” option, respectively. Dataset: MCF10A-1. (C) Visualization of identified cell lineage as in Figure 1F, but by EllipTrack with the “Global” option. (D) Comparison of the tracking performance between the “Density” and “Global” options. The highest score in each criterion is highlighted in bold green. Dataset: MCF10A-1. Scores of the “Density” option are reproduced from Table 1. (E) Relationship between the number of training samples and tracking performance. Down-sampling was performed by randomly selecting a subset of the original training datasets. Mean ± std (n=3) are shown for each down-sampling scheme. Scores that have no significant difference (p > 0.05, one-side t-test) from the “no down-sampling” scheme are highlighted in bold green. Scores without down-sampling are reproduced from Table 1. (F) Applicability of training datasets to tracking untrained movies. TRA score was not reported due to a lack of manual annotation. Scores with less than 10% difference are highlighted in bold green.

To account for this variable cell behavior, EllipTrack implements an option to perform time- and local cell density-dependent inference of cell migration speeds from the training datasets. Users can select the best option based on whether there exists a clear trend between the migration speeds and time/density values. As a demonstration, we analyzed the training datasets of the MCF10A movie and found that cells indeed migrated slower in the densely populated regions (Figure 2B, orange dots). Consistently, density-dependent inference (denoted as the “Density” option) revealed a clear dependency between migration distances and local cell densities, with cells moving four times faster in the sparsest regions compared to the densest ones (Figure 2B, black line). In comparison, assuming a constant migration speed over time (denoted as the “Global” option), inferred from the training datasets without the consideration of time or density values, does not reflect the distribution of cell behaviors, especially for the cells in the densely populated regions where their migration speeds were over-estimated by two-fold (Figure 2B, gray line).

To demonstrate how the Density option improves the tracking performance, we tracked the MCF10A-1 movie with both the Density (used in the previous benchmark) and Global options. Visualization of the same cell lineage in Figure 1F on CDK2 activity-based heatmaps revealed that the Density option tracked the entire lineage without errors (Figure 1F), whereas the Global option incorrectly swapped two cell tracks (Figure 2C). Furthermore, we systematically benchmarked these two options and found that the Density option outperformed the Global option in all track-linking criteria (Figure 2D). It is worth mentioning that even with the Global option, half of the identified complete tracks were error-free, which further demonstrates the power of the local track-correction module. In summary, the time- and density-dependent inference option complements the local track-correction module and allows users to achieve an even better tracking performance for appropriate movies.

### EllipTrack Requires Minimal Training

EllipTrack relies on user-provided training samples to construct cell tracks. A training sample describes a cell behavior (e.g. cell overlaps, mitosis, apoptosis, migration between two frames). When benchmarking the MCF10A-1 and A375 movies, we constructed 2922 and 2136 samples for cell behaviors, and labeled 37 and 55 cells for cell migration. To determine how many samples EllipTrack requires for accurate cell tracking, we down-sampled the training datasets to various degrees and examined the change in tracking performance. As shown in Figure 2E, EllipTrack displays a consistent performance when down-sampled five-fold, with no significant difference observed in any of the four track-linking criteria. With further down-sampling, tracking performance worsened and a marginally higher number of tracking errors occurred. This analysis suggests that creating ∼500 training samples and labeling ∼10 cells that are representative of cell behavior and migration are sufficient for high-accuracy tracking.

To determine whether training datasets from one movie could be used to track other movies, we created two additional movies of MCF10A (untreated) and A375 cells (treated with 1 μM dabrafenib) by imaging cells under identical conditions. We then compared the tracking performance using the previously generated training datasets vs. using training datasets generated from the new movies. In all four movies, EllipTrack demonstrated comparable tracking in both scenarios (Figure 2F), indicating good predictive power of the training datasets for tracking un-trained movies of the same cell line under the same imaging conditions. This brings substantial time savings since each cell line only needs to be trained once.

### EllipTrack Offers User-Friendly GUIs

The abundance of parameters and lack of intuitive interface often prevent users from learning new cell trackers (Ulman et al., 2017). To improve the practical usability of EllipTrack, we developed two user-friendly Graphical User Interfaces (GUIs), Parameter Generator GUI and Training Data GUI, to help users set up parameter values and construct training datasets (Figure S2). In Parameter Generator GUI, we classified parameters into two categories: basic and advanced. For most movies, setting the basic parameters is sufficient. We allow users to view the results of segmentation and migration-speed inference in real-time, thus providing a convenient parameter-tuning process. We also implement substantial hints and sanity-checks to prevent users from setting invalid values. In Training Data GUI, we simplify the training process such that creating a training sample involves only two mouse clicks and takes a few seconds. This allows users to generate the required ∼500 training samples within 1 hr. Together, we expect that these GUIs help users to become familiar with EllipTrack in a short amount of time.

## Discussion

Time-lapse microscopy provides an unprecedented opportunity to observe single-cell dynamics in real time. However, existing computational tools often track cells with limited accuracy and extensive human labor can be required to correct cell tracks manually, both of which prevent efficient high-throughput monitoring of cells over long time periods. This work describes EllipTrack, a new cell-tracking pipeline that combines the best features of previous segmentation and tracking algorithms and also implements a new local track-correction module to generate nearly error-free cell lineages. We also demonstrate EllipTrack’s ability to perform time- and density-dependent migration speed inference, characterize the amount of training data required for successful tracking, and introduce two user-friendly GUIs. With its superior performance, EllipTrack can be applied to address a wide range of biological questions. EllipTrack is especially suitable for drug screens and systems pharmacology, as it can handle large-scale movies with time-dependent changes in morphology and migration that occur upon drug treatment. EllipTrack has already proved essential in two projects from our lab, enabling new discoveries about proliferation in both normal and cancer cells (Min et al., 2020; Yang et al., 2020).

Despite its superior performance, EllipTrack has several limitations. First, due to the comprehensiveness of local track-correction, EllipTrack requires a significantly longer run time compared to other cell trackers, and it is often necessary to track multiple movies in parallel on a computing cluster. Second, cells with kidney-shaped or other non-elliptical nuclei are often over-segmented, although we find that signals can still be reliably extracted. Third, EllipTrack does not consider unusual cell behaviors such as multipolar mitoses and cell-cell fusion, although these behaviors can be incorporated by if needed. Similarly, whole-cell segmentation could also be incorporated via the use of whole-cell markers, if needed. In summary, EllipTrack outperforms all current state-of-the-art cell trackers, and the plethora of single-cell traces identified by EllipTrack can accelerate the discovery of heterogeneous cell behaviors over long time periods.

## AUTHOR CONTRIBUTIONS

C.T., C.Y. designed, conducted, and analyzed the experiments. C.T. developed and tested EllipTrack. S.L.S. conceived the project, suggested the experiments, and interpreted the data. C.T. and S.L.S. wrote the manuscript.

## ACKNOWLEDGEMENTS

We thank the members of the Spencer Lab for general help. This work was primarily supported by a Beckman Young Investigator Award to S.L.S. and in part by a Kimmel Scholar Award (SKF16-126), and a Searle Scholar Program Award (SSP-2016-1533) to S.L.S.

## SUPPLEMENTAL FIGURES

**Figure S1.**
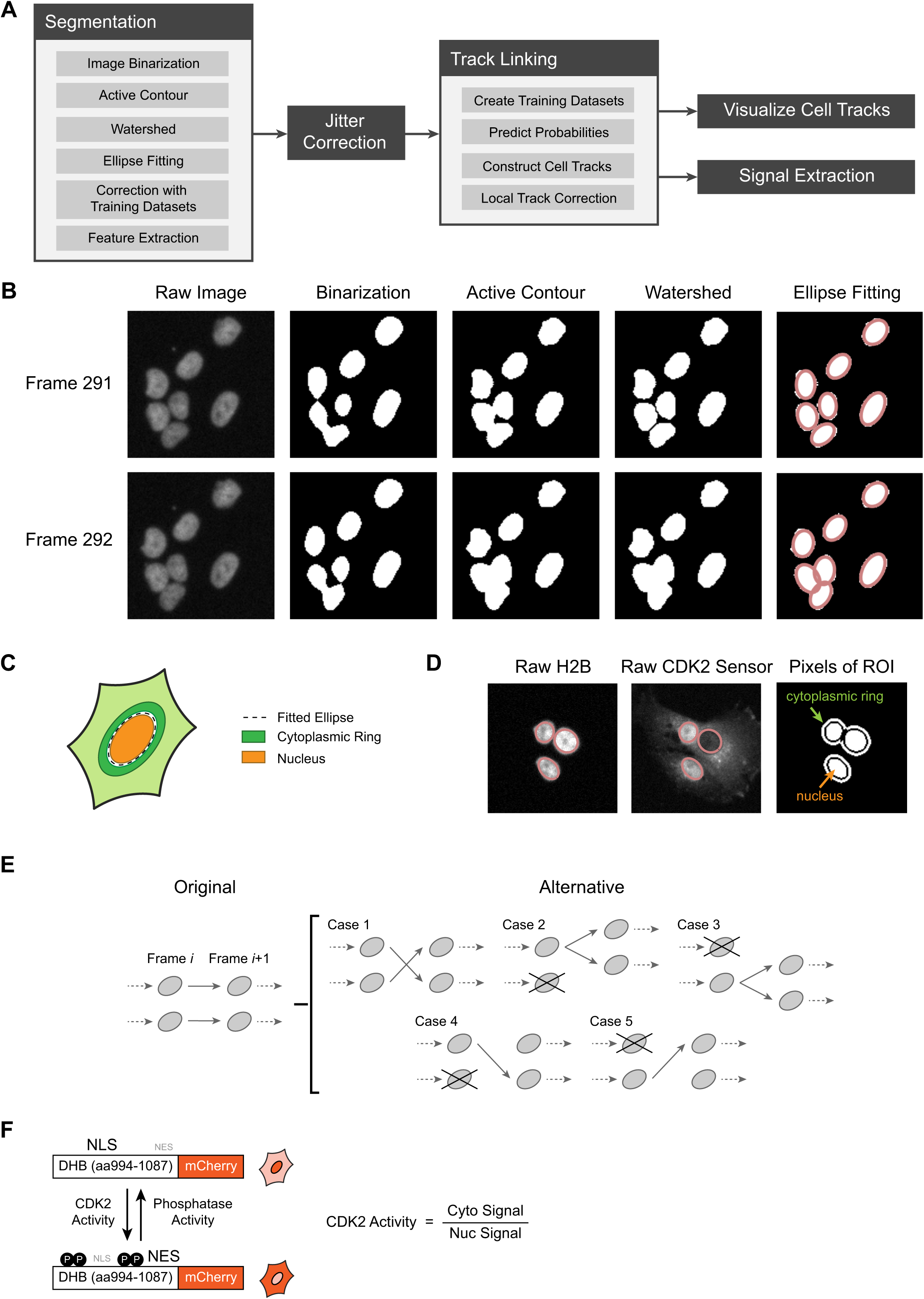
Implementation of EllipTrack, related to Figure 1. (A) Schematic illustration of the EllipTrack workflow. (B) Procedure of segmentation in EllipTrack. Raw images of the nuclear marker are first binarized. The binary images are then optimized with the Active Contour and Watershed algorithms. Finally, ellipses are fitted to the contours of the foreground pixels in the binary images. Dataset: A375-1. (C) Schematic illustration of the nuclear and cytoplasmic ring. (D) Visualization of the nucleus and cytoplasmic ring of overlapping nuclei. Left: Fitted ellipses plotted on top of the image of the nuclear marker (H2B). Middle: Fitted ellipses plotted on top of the CDK2 sensor image. Right: Pixels used to define the nuclear and cytoplasmic rings (white). Pixels shared by multiple cells are removed from the calculation. (E) Schematic illustration of the alternative track configurations. (F) Schematic illustration of the CDK2 sensor.

**Figure S2.**
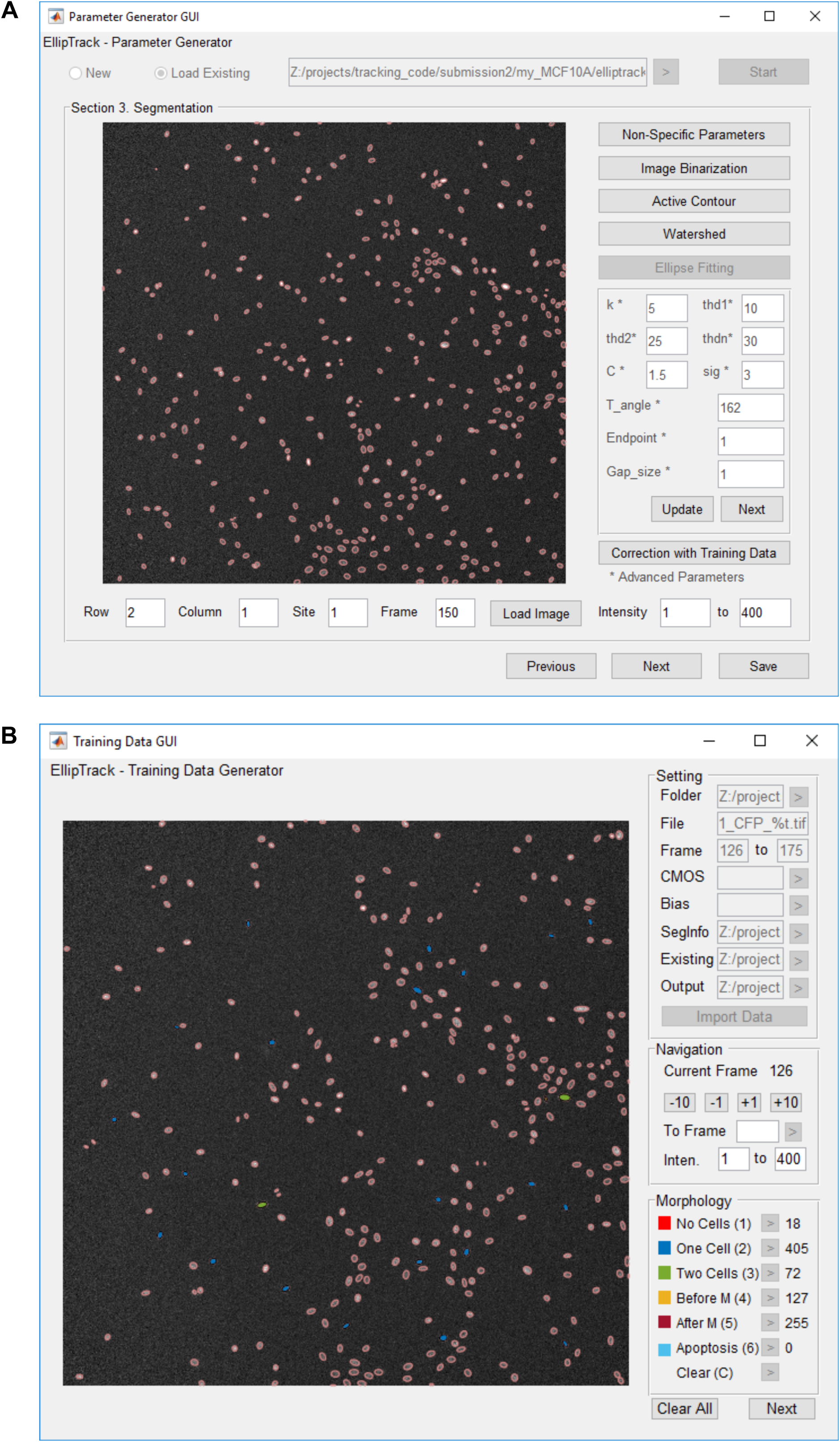
Graphical User Interfaces, related to Figure 1. (A) Parameter Generator GUI. (B) Training Data GUI.

## SUPPLEMENTAL TABLES

**Table S1.** Comparison of cell trackers, related to Table 1.

| Cell Tracker | Segmentation | Track Linking | Post-Processing |
| --- | --- | --- | --- |
| EllipTrack | Thresholding/blob detection, active contour, watershed, ellipse fitting | Viterbi algorithm | Local track correction |
| Baxter Algorithm | Bandpass filter, watershed | Viterbi algorithm | Post-tracking segmentation |
| Cappell *et al.* | Thresholding/blob detection, watershed | Nearest neighbor | / |
| iLastik | Random forest classifiers | Flow-based algorithm | Post-tracking segmentation |

**Table S2.** Statistics of benchmarked movies, related to Table 1. A detection refers to one nucleus in one frame. “% Detection Annotates” indicates the percentage of detections that were manually annotated for constructing the ground truth cell lineages.

| Movie | # Frames | # Cells in First Frame | # Cells in Last Frame | # Total Detections | Interval (min) | Duration (hr) | % Detections Annotated |
| --- | --- | --- | --- | --- | --- | --- | --- |
| HeLa-1 | 92 | 43 | 137 | $8.6 \times 10^3$ | 30 | 46 | 100 |
| HeLa-2 | 92 | 125 | 363 | $2.54 \times 10^4$ | 30 | 46 | 100 |
| BJ5TA | 107 | 329 | 332 | $3.54 \times 10^4$ | 12 | 21.4 | 100 |
| MCF10A-1 | 288 | 156 | 851 | $1.17 \times 10^5$ | 10 | 48 | 65 |
| A375-1 | 482 | 50 | 318 | $9.9 \times 10^4$ | 15 | 120.5 | 100 |
| RPE-hTERT | 400 | 30 | 712 | $1.05 \times 10^5$ | 15 | 100 | 64 |

**Table S3.** List of features, related to Figure 1.

| Category | ID | Description | Remark |
| --- | --- | --- | --- |
| Ellipse | 1 | Area | $\pi ab$ |
| Geometry | 2 | Major axis | Denote as $a$ |
| | 3 | Minor axis | Denote as $b$ |
| | 4 | Eccentricity | $\sqrt{a^2 - b^2}/a$ |
| | 5 | Equivalent radius | Denote as $r = \sqrt{ab}$ |
| | 6 | Approximate perimeter | $\pi\sqrt{2(a^2 + b^2)}$ |
| | 7 | Adjusted radius | $ab/\sqrt{a^2 + b^2}$ |
| Intensity of | 8 | Mean | Denote as $\mu_i$ |
| Ellipse Interior | 9 | Standard deviation | Denote as $\sigma_i$ |
| | 10 | Kurtosis | Denote as $\kappa_i$ |
| | 11 | Median | Denote as $m_i$ |
| | 12 | 75% percentile | Denote as $q_i$ |
| Intensity of | 13 | Mean | Denote as $\mu_b$ |
| Ellipse | 14 | Standard deviation |  |
| Boundary | 15 | Median |  |
| | 16 | Ratio of internal to boundary intensity | $\mu_i/\mu_b$ |
| | 17 | Approximate gradient | $(\mu_i - \mu_b)/r$ |
| Intensity of | 18 | Intensity of ellipse centroid | Denote as $\mu_c$ |
| Ellipse Centroid | 19 | Ratio of centroid to boundary intensity | $\mu_c/\mu_b$ |
| | 20 | Approximate gradient | $(\mu_c - \mu_b)/r$ |
| | 21 | Ratio of centroid to interior intensity | $\mu_c/\mu_i$ |
| | 22 | Approximate gradient | $(\mu_c - \mu_i)/r$ |
| Intensity Ratios | 23 | Standard deviation / Mean | $\sigma_i/\mu_i$ |
| of Ellipse | 24 | Kurtosis / Mean | $\kappa_i/\mu_i$ |
| Interior | 25 | Median / Mean | $m_i/\mu_i$ |
| | 26 | 75% percentile / Mean | $q_i/\mu_i$ |

## SUPPLEMENTAL MOVIE

**Movie S1. Comparison of cell trackers on tracking the MCF10A-1 dataset, related to Table 1.**

Tracking performance is visualized as described in Figure 1D. A cropped region of the movie consisting of 700×700 pixels (∼12% of the movie) is shown. This region does not include the image borders where iLastik frequently made tracking mistakes. IDs of cells whose centroid is located within the cropped region are displayed. Cell tracks were manually examined and tracking mistakes are visualized by orange arrows. A tracking mistake includes the four types illustrated in Figures 1D and 1F as well as incorrect track initiation, but does not include errors only due to segmentation. The total number of mistakes up to the current frame is displayed in the title of each panel. EllipTrack after local track correction out-performs the other cell tracks by making 2X to 9X fewer mistakes.

## KEY RESOURCE TABLE

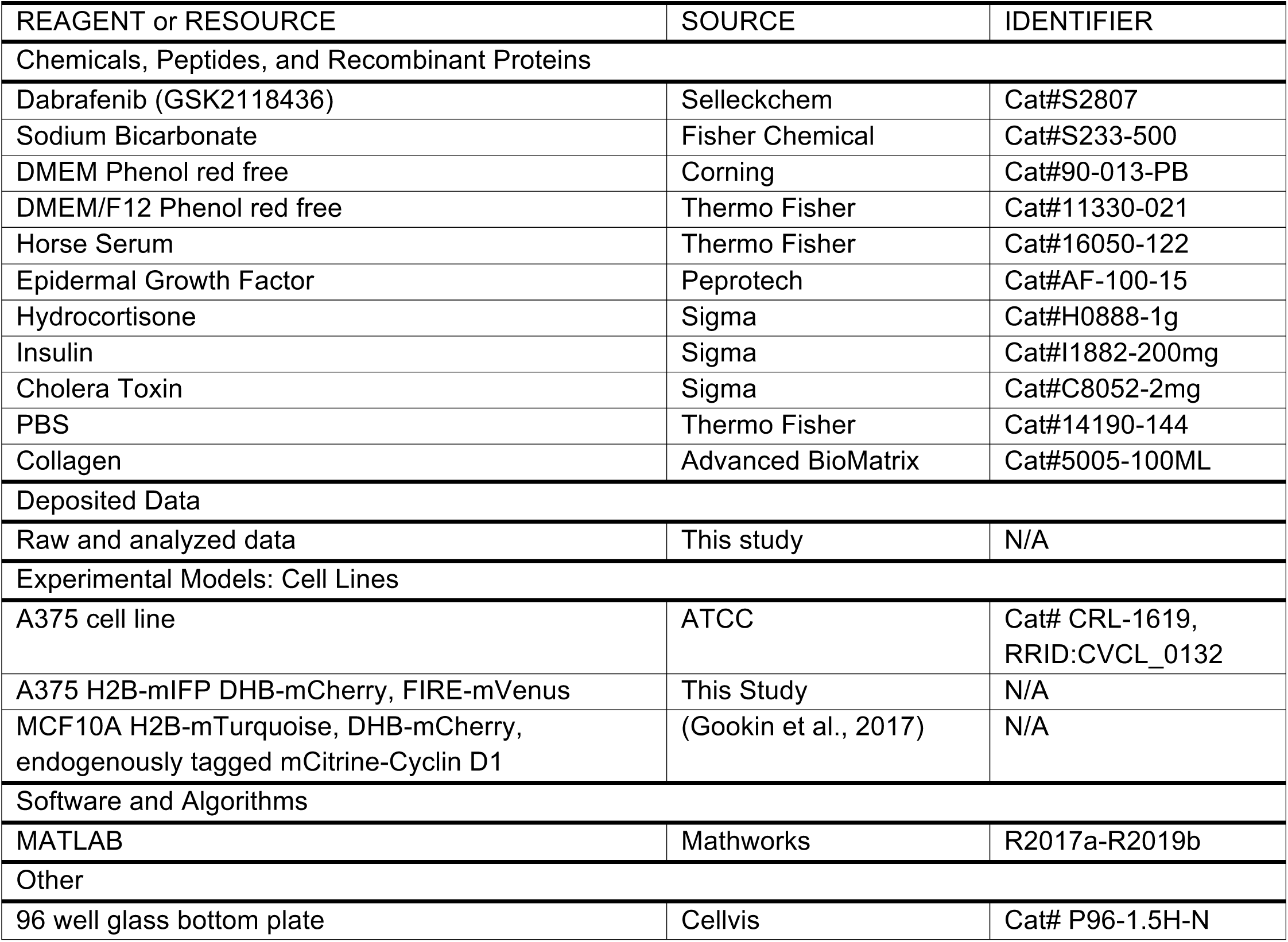

## METHODS

### Implementation of EllipTrack

A schematic diagram of EllipTrack is shown in Figure S1A. In brief, EllipTrack first segments the cell nuclei in the images of the nuclear marker and fits them with ellipses (Segmentation). Then, displacement between every two images in neighboring frames (called “jitters”) is calculated and the positional information of ellipses are corrected (Jitter Correction). Next, EllipTrack constructs cell tracks based on the ellipse information (Track Linking). Tracking results can be optionally visualized by creating the “vistrack” movies (Visualize Tracking). Finally, EllipTrack extracts signals from each ellipse and constructs time series for each cell track (Signal Extraction).

#### Segmentation

In Segmentation, an image of the nuclear channel is first background subtracted and binarized (“Image Binarization”) such that pixels within a nucleus are assigned a value of 1 (foreground pixels) and pixels in the image background are assigned a value of 0 (background pixels). The binary image is then optimized by the active contour (“Active Contour”) and Watershed (“Watershed”) algorithms. Next, ellipses are fit to the contours of foreground pixels in the optimized binary image (“Ellipse Fitting”) and corrected with user-provided training datasets (“Correction with Training Datasets”). Finally, EllipTrack extracts features from each ellipse that will be used in Track Linking (“Feature Extraction”). “Image Binarization”, “Ellipse Fitting”, and “Feature Extraction” are required, while the other three are optional.

In Image Binarization, two options are provided to binarize images: thresholding and blob detection. For thresholding, a threshold is applied to the intensities of the nuclear image. Pixels with intensities greater than the threshold are assigned as foreground pixels while other pixels are assigned as background pixels. For blob detection, the hessian matrix of the nuclear image is computed and a threshold is applied to the hessian matrix. Pixels with values less than the threshold are assigned as foreground pixels and other pixels are assigned as background pixels. A set of connected foreground pixels is called a component.

In Active Contour, the exact boundaries of components in the binary image are searched with the active contour algorithm. EllipTrack provides two options. The “global” option applies the active contour algorithm to the entire image, while the “local” option applies the algorithm to the local neighborhood of every component and the outputs from each local neighborhood are then assembled. The “global” option is computationally efficient, while the “local” option detects the boundaries more accurately, especially for dim nuclei.

In Watershed, multiple nuclei within a component are separated with the Watershed algorithm. To improve the robustness of the algorithm, the “peaks” of watershed are not defined by the pixels with local maximal distances to the background. Instead, they are defined by small components constructed by repeatedly eroding the binary image.

In Ellipse Fitting, contours of components in the binary image are fit with ellipses with a previously published algorithm (Zafari et al., 2015). In brief, the algorithm first searches for a characteristic point (called “seedpoint”) for each nucleus. The contours of components are then assigned to the seedpoints. Finally, the contour assigned to each seedpoint is fit with an ellipse.

In Correction with Training Datasets, EllipTrack constructs a linear discriminant classifier with user-provided training datasets (see “Construct Training Data” of “Track Linking”) and predicts the number of nuclei each ellipse contains. Ellipses with a high probability of containing no nucleus are removed, while ellipses with a high probability of containing two or more nuclei are split into two using the k-means algorithm. To ensure the robustness of splitting, two runs of the k-means algorithm are performed and the ellipse is only split if these two runs show a high level of agreement.

Finally in Feature Extraction, a predefined set of 26 features is extracted from each ellipse (Table S3). Features include the morphological parameters of the fitted ellipses and the distribution of nuclear marker intensities within the ellipses.

#### Jitter Correction

EllipTrack offers two options to perform jitter correction: local and global. With the local option, EllipTrack computes the jitters between every two neighboring frames (called “local jitters”) by image registration (as in (Cappell et al., 2016)), and then corrects the positional information of ellipses by subtracting the jitters from their original positions on the images. Movies are corrected independently, even if they are captured on the same plate.

Meanwhile, for movies captured on the same multi-well plate, jitters are often correlated because they are subject to the same plate motion. It is therefore possible to infer the plate motion from this correlation and use it to compute the jitters for each movie based on its position on the plate, an approach that can significantly reduce the errors of calculation. EllipTrack implements this approach with the global option, where the plate motion between every two frames is inferred from the local jitters, and the jitters calculated from the plate motion (called “global jitters”) are then used to correct the positions of ellipses.

Mathematically: denote the position of well (*i, j*) in frame *t* as (*x*_*i,j*_^*t*^, *y*_*i,j*_^*t*^) and the distances between two rows and two columns as *v* and *h*, respectively, as below

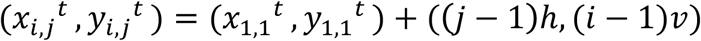

Assume that the plate motion between frame *t* and frame *t*+1 causes a whole-plate shift of (*Δx, Δy*) and a plate rotation of angle *θ*. The position of well (*i, j*) in frame *t*+1 can then be expressed as

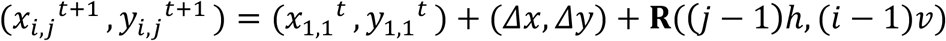

where **R** is the rotation matrix for the angle *θ*. Therefore, the motion of the well (i.e. global jitters) follows

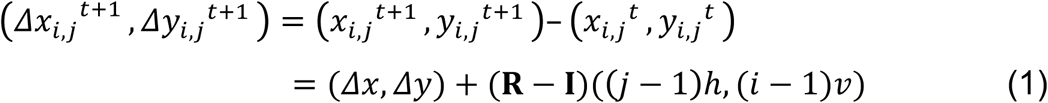

with **I** being the identity matrix. Local jitters inferred by image registration are equal to the sum of the global jitters and the error terms, as shown below.

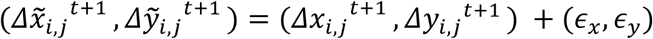, where *ϵ*_I_ and *ϵ*_J_ ∼ *N*(0, 1) EllipTrack uses the local jitters 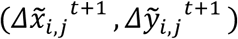 and the position (*i, j*) of each well to infer the unknown variables (*h, v, θ, Δx, Δy*) and then computes the global jitters of each well with Equation 1.

#### Track Linking

In Track Linking, users first create training datasets with the Training Data Generator GUI attached to EllipTrack (“Construct Training Data”) and EllipTrack uses these datasets to predict the behaviors of each ellipse in the movie (“Compute Probabilities”). Cell tracks are then generated based on these predictions (“Construct Tracks”). Finally, errors in cell tracks are fixed by the local track-correction module (“Local Track Correction”).

In Construct Training Data, users load ellipse information into the GUI attached to EllipTrack and manually label the behavior (called “events”) of each ellipse. Two types of events are considered: morphological events and motion events. Morphological events describe the status and behavior of the ellipse in the current frame and include the number of cell nuclei in the ellipse (0, 1, or 2) as well as whether cells are mitotic, newly born, or apoptotic. Motion events describe cell migration between neighboring frames and include migration between two ellipses in neighboring frames and migration in/out of the field of view. Motion events are also used to infer the migration speed of cells. After users label all events of interest, the GUI generates a training dataset which can be used for computing probabilities.

In Compute Probabilities, EllipTrack constructs linear discriminant classifiers with user-provided training datasets and computes the probabilities of morphological events for each ellipse. For motion events, EllipTrack models cell migration by random walk (as in (Magnusson et al., 2015)), and calculates the probabilities of cell migration with the formula

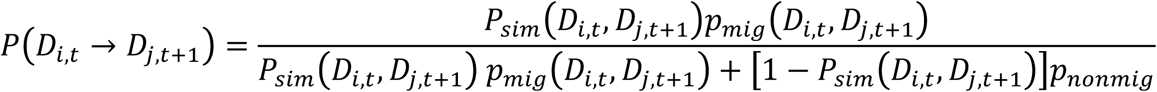

Here, *D*_*i,t*_ and *D*_*j,t*+1_ refers to the *i*-th ellipse in Frame *t* and *j*-th ellipse in Frame *t*+1; *P*_*sim*_ *D*_*i,t*_, *D*_*j,t*+1_ refers to the probability that these two ellipses represent the same cell, as predicted by the linear discriminant classifiers constructed from the training datasets; *p*_*mig*_ *D*_*i,t*_, *D*_*j,t*+1_ refers to the probability that the cell migrates from the position of *D*_*i,t*_ to the position of *D*_*j,t*+1_, as computed by the formula of random walk; and *p*_*nonmig*_ refers to the null probability of cell migration (a user-controlled parameter). A key parameter of a random walk is the standard deviation (i.e. cell migration speed). EllipTrack offers three options to compute this parameter: global, time, and density. With the global option, EllipTrack uses all training samples and infers a single migration speed for all cells in all frames. With the other two options, EllipTrack calculates the migration distances and the Frame ID (time) or the number of cells in the local neighborhood (density) for each training sample, and estimates the curve of migration speed vs time/density from these samples. EllipTrack will then apply this curve to the movie and calculate the migration speed for each cell at each frame.

In Construct Tracks, EllipTrack uses the Magnusson global track-linking algorithm (Magnusson et al., 2015) to construct cell tracks. Note that in EllipTrack, the Magnusson swap operation only applies to the cell tracks that map to cells at both frames. Cell tracks that map to cells at only one frame at the frames of interest, such as those associated with appearance, disappearance, mitosis, and apoptosis, are not considered. In comparison, all cell tracks can be swapped in BaxterAlgorithm. Therefore, BaxterAlgorithm may show a superior performance compared to EllipTrack before local track-correction (Table 1). However, all these overlooked mistakes can be fixed by the local track-correction module, and the tracking performance of EllipTrack after correction often becomes much better than BaxterAlgorithm.

Finally, in Local Track Correction, EllipTrack employs a six-step procedure to correct possible tracking mistakes. In the first step, EllipTrack optimizes the Magnusson swap operation and examines alternative track configurations, as described in the main text. In the second step, EllipTrack corrects premature termination of cell tracks due to under- and over-segmentation. In the first case, when two cells are under-segmented by one ellipse and the predicted probabilities of cell overlap are not perfect, one cell might lose its cell track at the frame of under-segmentation and later be mapped by a new cell track at the frame where segmentation becomes correct. To fix this type of mistake, EllipTrack examines every pair of prematurely terminated cell track and newly created one, and will connect them if they meet the following requirements: (1) the latter cell track appears at most ‘track_para.critical_length’ (a user-controlled parameter) frames later than the termination of the former cell track, (2) both cell tracks share the same cell as their closest neighbor, and (3) the latter cell track does not appear by a mitosis event. In the second case, when one cell is over-segmented by two ellipses, its cell track might map to one of the two ellipses and terminates, while another cell track appears, maps to the other ellipse, and continues mapping this cell after segmentation becomes correct. To fix this type of mistake, EllipTrack examines every pair of prematurely terminated cell track and newly created one, and will combine them if they meet the following requirements: (1) the latter cell track appears at most ‘track_para.critical_length’ frames earlier than the termination of the former cell track, (2) both cell tracks are the closest neighbor of each other, and (3) the former cell track does not terminate by a mitosis event.

In the third step, EllipTrack corrects mistakes associated with under-segmentation. Due to its memoryless nature, the Magnusson global track-linking algorithm loses the identities of the cells when two cells are under-segmented and both cell tracks map to the same ellipse. Consequently, the algorithm randomly maps the cell tracks when cells are no longer under-segmented, which results in 50% of such mappings being incorrect. To correct these mistakes, EllipTrack compares the ellipse similarity and migration distances before and after under-segmentation and swaps the two cell tracks after under-segmentation if both quantities improve.

In the fourth step, EllipTrack searches for undetected mitosis events, which are marked by a cell track mapping both the mother cell and one daughter cell, and a second cell track mapping the other daughter cell. To do so, EllipTrack looks for all cell tracks which are not created by mitosis but have high probabilities of being a newly born cell at their first frame (as predicted in “Compute Probabilities”). For each cell track, EllipTrack then examines whether there exists a neighboring cell track that has a high probability of being a mitotic cell in the previous frame and has a high probability of being a newly born cell in the current frame. If such a cell track is found, EllipTrack will split the mother-and-daughter cell track into two (one for the mother cell, and the other for the daughter cell) and create a mitosis event between the mother cell track and the two daughter cell tracks.

In the fifth step, EllipTrack prunes the cell lineage trees by removing all cell tracks unlikely to represent a cell, such as very short cell tracks and cell tracks skipping too many frames. Mitosis events associated with these invalid cell tracks will be removed as well.

In the last step, EllipTrack recursively adjusts the Track IDs such that (1) Track IDs will be sorted by the ellipse positions at the first frame, and (2) sibling cells (two cells born by the same mother) have consecutive Track IDs. This step significantly reduces the mental burden when examining the “vistrack” movies.

#### Visualize Tracking

To visualize the cell tracks, EllipTrack generates “vistrack” movies as in Figure 1D.

#### Signal Extraction

In Signal Extraction, EllipTrack defines the regions of nucleus and cytoplasmic ring for each ellipse as illustrated in Figure S1C and removes a pixel from the region if it is shared by other regions or cells (Figure S1D). EllipTrack then performs background subtraction on each fluorescent image, drops the outlier intensities (defined as the top and bottom 5%), and computes the mean, median, and variance of signal intensities in each region of each ellipse. Finally, the signal time series in each cell track are assembled for plotting.

### Generation of MCF10A and A375 Movies

Time-lapse microscopy was performed as previously described (Arora et al., 2017). In brief, cells were plated on a 96-well plate coated with collagen (1:50 dilution in water) in phenol red-free full growth medium. Approximately 24 hr later, cells were transferred to a Ti-E PFS (Nikon) with a humidified, 37°C chamber with 5% CO_2_, and imaged periodically with a 10X 0.45 NA objective. Drug was added to the medium by pausing the movie, exchanging 50% of the medium in the well with medium containing 2X drug concentration, and restarting the movie. Movie-specific parameters are described below.

#### A375 Movie

A375 cells expressing H2B-mIFP (Yu et al., 2015), DHB-mCherry, and FIRE-mVenus (Albeck et al., 2013) were plated with a density of 1000 cells/well. Images were taken every 15 min with the following exposure times: mIFP 300 ms, mCherry 50 ms, and YFP 100 ms. Data from the mCherry and YFP channels was not used in this study. Cells were first imaged in full growth medium (DMEM supplemented with 10% FBS, 1.5 g/L sodium bicarbonate, and 1X penicillin/streptomycin) for 24 hr (96 frames). The BRAF inhibitor dabrafenib was then added to the well as described above and cells were imaged for a further 96.5 hr (386 frames). Drug was refreshed 30 hr after drug addition.

#### MCF10A Movie

MCF10A cells expressing H2B-mTurquoise, DHB-mCherry, and endogenously tagged mCitrine-Cyclin D1 (Gookin et al., 2017) were plated with a density of 1000 cells/well and cultured in full growth medium (DMEM/F12 supplemented with 5% horse serum, 100 ng/mL cholera toxin, 20 ng/mL epidermal growth factor, 10 μg/mL insulin, 0.5 mg/mL hydrocortisone, and 1X penicillin/streptomycin). Cells were imaged every 10 min for 48 hr (288 frames) with the following exposure times: CFP 20 ms, and mCherry 100 ms. The YFP channel was not imaged.

### Description of Movies

Movie statistics were summarized in Table S2. A detection refers to one nucleus in one frame. Based on the total number of detections, we classified MCF10A, A375, and RPE-hTERT movies as large-scale.

#### HeLa

Training movies of the Fluo-N2DL-HeLa dataset from the Cell Tracking Challenge. In each movie, HeLa cells expressing H2B-GFP were imaged every 30 min for 46 hr (92 frames) (Maška et al., 2014; Ulman et al., 2017). Official ground truth was used. These movies were among the most challenging datasets from Cell Tracking Challenge.

#### BJ5TA

Courtesy of Steven Cappell. BJ5TA cells expressing H2B-mTurquoise and FUCCI-mCherry (Sakaue-Sawano et al., 2008) were treated with control siRNA and then imaged every 12 min for ∼20 hr (107 frames) (Cappell et al., 2016). Movie jitters were not corrected. 800 cell nuclei from four frames (Frame 23, 44, 81, 105) were annotated for segmentation benchmark and all cell lineages were annotated for tracking benchmark. The main difficulties include a drastic change of image background due to photobleaching, strong illumination bias, heterogeneous brightness of nuclei, and interference from other channels.

#### MCF10A

This movie was generated in this study. Experimental details were described above. Movie jitters were not corrected. 802 cells from four frames (Frame 41, 133, 217, 273) and all the cell lineages that present in the first frame were annotated. The main difficulties include rapid cell migration and dense cell distribution.

#### A375

This movie was generated in this study. Experimental details were described above. Movie jitters were corrected by BaxterAlgorithm. 668 cells from six frames (Frame 57, 106, 201, 210, 318, 480) and all cell lineages were annotated. The main difficulties include a drastic change of image background due to photobleaching, as well as abnormal cell morphology and strong overlap after drug treatment. Best efforts were made to annotate overlapping cells, though multiple cell lineages were not annotated to the end of the movie due to severe overlapping.

#### RPE-hTERT

Courtesy of Tammy Riklin Raviv and Jose Reyes. RPE-hTERT cells expressing H2B-mTurquoise, DDX5-eYFP, and p21-mVenus were imaged every 15 min for 100 hr (400 frames). Medium was refreshed at Frame 92, 181, 282, and 374 (Arbelle et al., 2018). Movie jitters were not corrected. For segmentation benchmark, manual annotation from Arbelle *et al.* were used (502 cells, from Frame 1, 20, 40, 60, 61, 80, 100, 190). For tracking benchmark, additional annotation was provided such that the majority of cell lineages, including all lineages that present in the first frame, were annotated. Main difficulties include strong cell overlap, rapid cell migration and dense cell distribution.

### Benchmarking Criteria

#### Cell Tracking Challenge Criteria (SEG and TRA)

These criteria evaluate the resemblance between tracking results and manually annotated ground truth. Ground truth was obtained as described above. Official evaluation programs were used to compute the scores. Since EllipTrack and Cappell *et al.* do not perform post-tracking segmentation, multiple cell tracks might map to the same nuclei, which is incompatible with the official evaluation programs. We therefore partitioned all such under-segmented nuclei with k-means algorithm and assigned track IDs based on the positions of nuclear centroids.

#### Accuracy of Segmentation (%CORR_S)

This criterion examines whether cell nuclei are accurately segmented, without post-tracking segmentation. All cells from five equidistant frames spanning over the entire movie were examined (> 500 cells). Every cell nucleus was manually classified into four categories: correctly segmented, under-segmented (contains more than one nucleus), over-segmented (contains less than one nucleus), and not segmented. The percentage of nuclei that were correctly segmented was reported.

#### Number of Complete Tracks (#COMP)

This criterion examines the ability of cell trackers to track cells over long time periods. A complete track refers to a cell being tracked along the lineage tree from the first to the last frame of the movie. Cells migrating in/out of the field of view are excluded, regardless of whether they are correctly tracked or not. Ground truth of #COMP was derived from the manually annotated cell lineages.

#### Accuracy of Complete Tracks (#MIS_T and %CORR_T)

These criteria examine the ability of cell trackers to generate error-free cell tracks. Every complete track was manually examined and all tracking mistakes were annotated. A tracking mistake includes incorrect mapping between frames, incorrectly assigned mitosis events, and undetected mitosis events, but does not include mistakes due to segmentation. The average number of tracking mistakes in the complete tracks was reported as #MIS_T, and the percentage of complete tracks that have no tracking mistakes was reported as %CORR_T.

### Parameters used for Benchmarking

Parameter files of EllipTrack, BaxterAlgorithm, and Cappell *et al.* can be accessed at https://drive.google.com/drive/folders/1dEW-cDIwZLrgA-yPLljuB2Ok2DHV2qQS. iLastik files are available upon request.

#### EllipTrack

For segmentation, parameters were tuned in Parameter Generator GUI such that cells from all frames were generally well-segmented. In brief, images were first background subtracted (HeLa only), log-transformed, and binarized (MCF10A and RPE-hTERT: thresholding, others: blob detection). Components in the binary images were then optimized with Active Contour (all but A375: log-transforming the raw images, all with the local option) and partitioned with Watershed. Finally, ellipse fitting was performed to the boundaries of components with the default parameters. For Track Linking, density-dependent migration speed inference was used for MCF10A and RPE-hTERT, while the global option was used for other movies. Default parameter values were used, except HeLa and A375 movies where some advanced parameters were tuned to improve the detection of mitosis events.

#### BaxterAlgorithm

Parameters provided in BaxterAlgorithm were used to track the HeLa movies. For all other movies, the HeLa-movie parameters were imported and their values were then modified as follows. For Segmentation, the parameter ‘BPSegThreshold’ was fine-tuned to achieve a good segmentation performance. Additional parameters, such as ‘SegMinArea’, might also be tuned to remove invalid components. For Track Linking, ‘pDeath’, ‘TrackPAppear’, ‘TrackPDisappear’, and ‘TrackNumNeighbours’ were modified to the same values as EllipTrack, and ‘TrackXSpeedStd’ was modified to the inferred migration speed by EllipTrack with the Global option. ‘pCnt0’, ‘pCnt1’, and ‘pCnt2’ were set to approximately match the proportions of over-, correctly, and under-segmented cells. Three parameters were tested with multiple values: ‘TrackMaxMigScore’ (0 or Inf), TrackMigLogLikeList (‘MigLogLikeList_uniformClutter’, or ‘MigLogLikeList_viterbiPaper’), and count/split/death classifiers (‘none’ or the ones created by training a subset of movie images). Tracking results from all eight parameter combinations were evaluated by the Cell Tracking Challenge evaluation software, and the combination with the highest (SEG+TRA)/2 score was used for benchmark.

#### Cappell *et al*

Parameters related to movie specification, such as magnification of objective lens and average nuclear radius, were modified to match the experiments. Three parameter combinations were tested based on how the first and later frames were segmented: (1) the first frame with the ‘log’ option while later frames with the ‘apriori’ option; (2) the first frame with the ‘single’ option while later frames with the ‘apriori’ option; and (3) all frames with the ‘log’ option. The parameter combination with the highest (SEG+TRA)/2 score was used for benchmark.

#### iLastik

For Segmentation, around 100 typical cells were selected from the entire movie. Pixels within the cell nuclei were trained as “Label 1” while the pixels immediately surrounding the nuclear contours were trained as “Label 2”. Additionally, pixels from image background were trained as “Label 2”. All features with the default standard deviation values were used for training. For Tracking, probabilities of segmentation were first smoothed with a sigma of 2×2, and then binarized with a threshold of 0.5. Components with area less than 25 (MCF10A) or 100 pixels (all other movies) were removed. Next, division and object count classifiers were trained on the same cells as EllipTrack. Training samples might be added or removed to avoid assertion errors. All features except locations were used for training. Finally, movies were tracked with the default parameters. Three values of border widths (HeLa, RPE-hTERT: 25, 50, 100 pixels, others: 50, 100, 200 pixels) were tested for each movie and the one with the highest (SEG+TRA)/2 score was used for benchmark.

### Availability of EllipTrack

EllipTrack is available at https://github.com/tianchengzhe/EllipTrack. Documentation is available at https://elliptrack.readthedocs.io/en/latest/. Movies and scripts used in this study are available upon request.

